# Anatomical and biophysical characterization of intergeneric graft-incompatibility within the Solanoideae

**DOI:** 10.1101/2022.07.01.498506

**Authors:** Hannah R Thomas, Alice Gevorgyan, Margaret H Frank

## Abstract

Interspecies grafting is a technique that allows beneficial shoot and root combinations from separate species to be combined into a single organism. Despite its relevance to agricultural production, little is known about the determinants of graft compatibility. One hypothesis for compatibility revolves around the taxonomic degree of relatedness between the two plants. To test how phylogenetic distance affects interspecific graft compatibility within the economically important Solanaceae sub-family, Solanoideae, we characterized the anatomical and biophysical integrity of graft junctions for graft combinations made between four species: tomato (*Solanum lycopersicum*), eggplant (*Solanum melongena*), pepper (*Capsicum annuum*), and groundcherry (*Physalis pubescens*). We analyzed the survival, growth, and junction integrity via bend tests, and imaged the cellular composition of the graft junctions in order to deduce the status of vascular connectivity across the junction. Utilizing these techniques, we were able to quantitatively assess the degree to which each interspecific combination exhibits compatibility. Despite the fact that most of our graft combinations exhibited high survival rates, we show that only tomato and eggplant heterografts are truly compatible. Unlike incompatible grafts, the formation of reconnected vascular tissue within the tomato and eggplant heterografts contributed to biophysically stable grafts that were resistant to snapping. Furthermore, we identified 10 graft combinations that show delayed incompatibility, providing a useful, economically relevant system to pursue deeper work into genetic and genomic determinants of graft compatibility. This work provides new evidence indicating that graft compatibility may be limited to intrageneric combinations within the Solanoideae subfamily. Further research using more extensive graft combinations amongst Solanaceous species can be used to test whether our hypothesis broadly applies to this family.

## 1. INTRODUCTION

Grafting is an ancient agricultural practice, where distinct plant parts are combined into a single organism (Mudge et al., 2009a). The root system is known as the stock and the grafted vegetative portion is known as the scion (indicated as Scion:Stock throughout this paper). The term graft-compatibility refers to the capacity for a given scion and stock combination to regenerate and stably reconnect their non-vascular and vascular tissue within an anatomically unique region, referred to as the graft junction (Benda et al., 1960; Rasool et al., 2020). Despite our limited understanding regarding the determinants of graft compatibility, it has been noted for thousands of years that taxonomic relatedness is a good predictor of graft success (Andrews and Marquez 2010; Pease 1933; Goldschmidt 2014; Mudge et al. 2009)(Andrews and Marquez 2010). For example, intraspecific grafts are generally more likely to be compatible than interspecific grafts (Goldschmidt, 2014; Mudge et al., 2009b). In contrast, there are some families that exhibit wide graft compatibility, including many examples of intergeneric graft compatibility, such as Rosaceae where economically relevant species are routinely grafted together from seperate genera, such as apples (*Malus domestica*) and pears (*Pyrus communis*; Errea et al., 1994; Westwood, 1993).

The Solanaceae or nightshade family is another such family that is often regarded as having wide graft compatibility (Westwood, 1993; Zeist et al., 2018; Kawaguchi et al., 2008). Recent work within the nightshade family, highlights this broad compatibility, demonstrating that individuals from the *Nicotiana* and *Petunia* genera can be grafted with diverse species, including across family limits (Notaguchi et al. 2020; Kurotani et al. 2022). Despite the abundance of economically important plants contained within the Solanoideae sub-family, little is known about the limitations of interspecific graft compatibility within this group, with a few notable exceptions (i.e. - tomato:eggplant, eggplant:tomato, tomato:potato, and eggplant:potato combinations; Liu et al., 2009; Romano and Paratore, 2001; Thompson and Morgan). Previous work has shown that tomato and pepper are incompatible and fail to form vascular reconnections within the first week of grafting (Kawaguchi et al., 2008; Thomas et al., 2022). Despite this knowledge, the graft compatibility between pepper and other closely-related crops remains unknown (Kawaguchi et al., 2008; Zeist et al., 2018; Thomas et al., 2022).

To determine the taxonomic limits that constrain intrafamily graft compatibility, we conducted a graft trial with four Solanoideae crops: tomato (*Solanum lycopersicum var. M82*), eggplant (*Solanum melongena var. BARI-6*), pepper (*Capsicum annuum var. California Wonder*), and groundcherry (*Physalis pubescens*). All four species belong to the Solanoideae sub-family, which is estimated to have diverged from Nicotianoideae 24 million years ago (ma; Figure 1; Särkinen et al., 2013). *Solanum*, which contains over half of the species in Solanaceae, has an estimated divergence time from *Capsicum* and Physalinae 19 ma (Frodin, 2004; Stern and Bohs, 2012; Särkinen et al., 2013). Soon after this divergence, *Capsicum* split into separate subclades approximately 18 ma (Särkinen et al., 2013). Eggplant and tomato, both members of *Solanum*, are the most closely related of the four crops in our study and are estimated to have diverged 14 ma (Särkinen et al., 2013).

**Figure 1:**
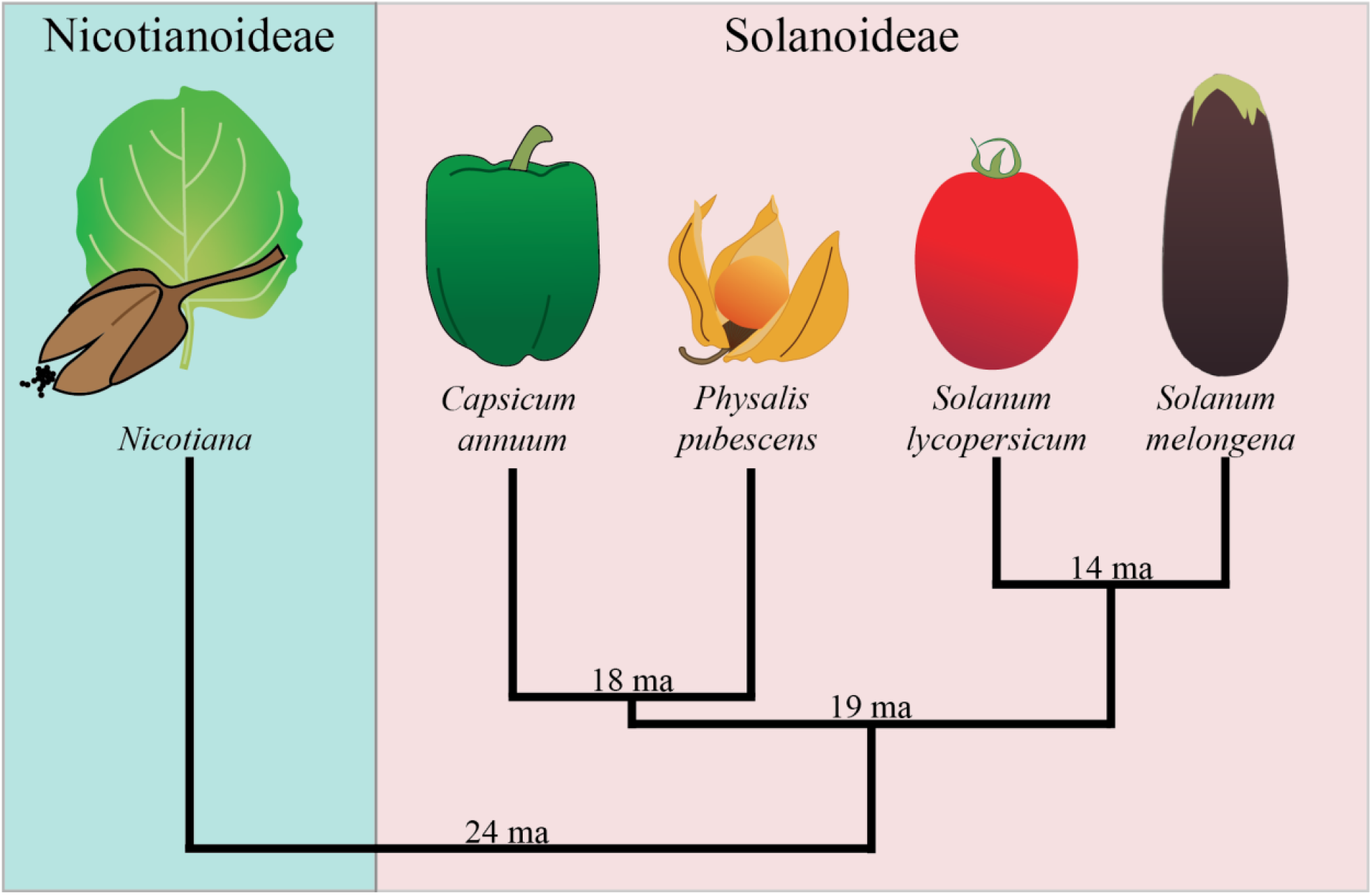
Phylogenetic relationship between Solanaceae sub-families Nicotianoideae and Solanoideae. Nicotianoideae and Solanoideae are sister sub-families within the Solanaceae. Three Solanoideae tribes are represented: Capsiceae (*Capsicum annuum*), Physaleae (*Physalis pubescens*), and Solaneae (*Solanum lycopersicum* and *Solanum melongena*). The estimated time of divergence is shown above the root of each branch (Särkinen et al., 2013).

Graft survival is often equated with graft compatibility. Our work within Solanoideae examines the deeper anatomical basis for compatibility and demonstrates that survival can occur in the absence of true compatibility. In order to test compatibility, we performed an extensive reciprocal graft trial, analyzed the vascular tissue in the graft junction, conducted stem stability tests, and measured lateral stem growth 30 days after grafting. By testing the graft junctions, we show that these four Solanoideae species, while broadly capable of surviving grafting, are generally not compatible, with the exception of intrageneric grafts between tomato and eggplant. To aid in the rapid identification of incompatible graft combinations, and avoid future conflation of survival and compatibility, we demonstrate that a simple bend test for biophysical stability within the junction can be used to diagnose delayed graft incompatibility.

## 2. MATERIALS AND METHODS

### 2.1 Plant materials and growth conditions

200 *Capsicum annuum* var. California Wonder and 200 *Physalis pubescens* seeds were stratified with 50% bleach for 30 seconds and then rinsed five times with sterile distilled water. The seeds were planted directly into LM-111 soil and kept on heat mats until grafted. 100 *Solanum lycopersicum* var. M82 and 80 *Solanum melongena* var. BARI-6 seeds were bleach-stratified and placed into Phyatrays© in the dark for 3 days, then moved to light for 3 days, and finally transferred into LM-111 soil on heat mats until grafted. All seedlings were grown in climate-controlled chambers set to 23 C with 16:8 day/night light cycles under F54T5/841/HO fluorescent bulbs (500-800 μmol/m^2^/sec).

### 2.2 Graft conditions

Four-week-old pepper and groundcherry seedlings, and two-week-old tomato and eggplant seedlings were used for grafting. This seedling germination timeline was optimized so the four species were of similar stem diameter (1.5-1.75 mm) and hypocotyl length. The following grafts were performed between tomato (T), pepper (P), groundcherry (GC), and eggplant (E): T:T, P:P, GC:GC, T:P, P:T, T:GC, GC:T, P:GC. GC:P (n = 20) and E:E, T:E, E:T, P:E, E:P, GC:E, and E:GC (N = 15). Any combination including eggplant only had 15 graft replicates due to the low availability of the BARI-6 seed variety. Scion and stocks were joined with a slant graft below the cotyledons (Kubota et al., 2008). Grafts were held together with 1.5 mm silicon-top grafting clips (Johnny’s Selected Seeds, Albion, Maine, USA). Grafted plants were generously watered, covered with plastic domes, and placed in the dark for 3 days. On day 4, plants were returned to light (500-800 μmol/m^2^/sec). To test Solanoideae graft compatibility, survival of each graft combination was noted on 0, 5, 7, 10, 14, 21, and 30 DAG. Herbaceous grafts heal within the first week (Melnyk, 2017); observing survival up to 30 DAG allowed us to track how survival changes over time in response to physiological properties like xylem connectivity.

### 2.3 Whole plant imaging and stem phenotyping

Seedlings were imaged prior to grafting, immediately following grafting, and 30 DAG using a 12-megapixel wide-angle camera (Samsung, South Korea). The shoots of 30 DAG plants were imaged in their pots. The stem of the seedlings prior to grafting and the scion and stock directly above or below the graft junction were measured 30 DAG using digital calipers.

### 2.4 Bend test

Graft junction integrity was tested using the bend test (Thomas et al., 2022). Due to the variable nature of graft survival, biological replicates vary (where T= tomato, P = pepper, GC = groundcherry, and E = eggplant): T:T n= 15, P:P n=9, E:E n=5, GC:GC n=12, T:P n=9, P:T n=7, T:E n=5, E:T n=9, T:GC n=6, GC:T n=6, P:GC n=1, GC:P n=8, P:E n=1 E:P n=9. GC:E and E:GC had too few replicates to test.

### 2.5 Tissue collection for confocal imaging

Graft junctions were harvested by cutting 2 cm above and below the cut site. Tissue was fixed and stained with propidium iodide as previously described (Thomas et al., 2022). Fully cleared graft junctions were imaged on a Zeiss LSM880 Confocal Microscope (Germany) using an Argon Laser 514 nm beam.

### 2.6 Statistical analysis of grafted plants

All statistical computations and graph generation were performed in R (R Core Team, 2021). Statistical significance of survival and stem integrity were calculated using Fisher’s exact test (Hervé 2020). All interval data were tested for normal distribution and homoscedasticity using Wilks-Shapiro test and Levene’s test from the CAR package (Fox and Weisberg, 2018). All data was found to be homoscedastic but non-parametric (Table S5, Figure S4). Analysis of variance (ANOVA) is robust enough to tolerate this violation. Tukey’s honest significant difference test with an adjusted p-value was used to determine pairwise significance. Plots were made in R using ggplot2 and dplyr (R Core Team 2020, Wickham 2011).

## 3. RESULTS

### 3.1 Solanoideae inter-species grafts have varying survival rates 30 Days After Grafting (DAG)

We selected four Solanoideae crops (tomato, eggplant, pepper, and groundcherry) from three different tribes (Capsiceae, Physaleae, and Solaneae) to test the taxonomic limits of graft compatibility. These four species were chosen because of their economic importance, as well as their suitability as graft partners based on their similar stem compositions, stem diameters, and growth rates (Figure 2, Figure S1, Table S1-, see Supplemental Data).

**Figure 2:**
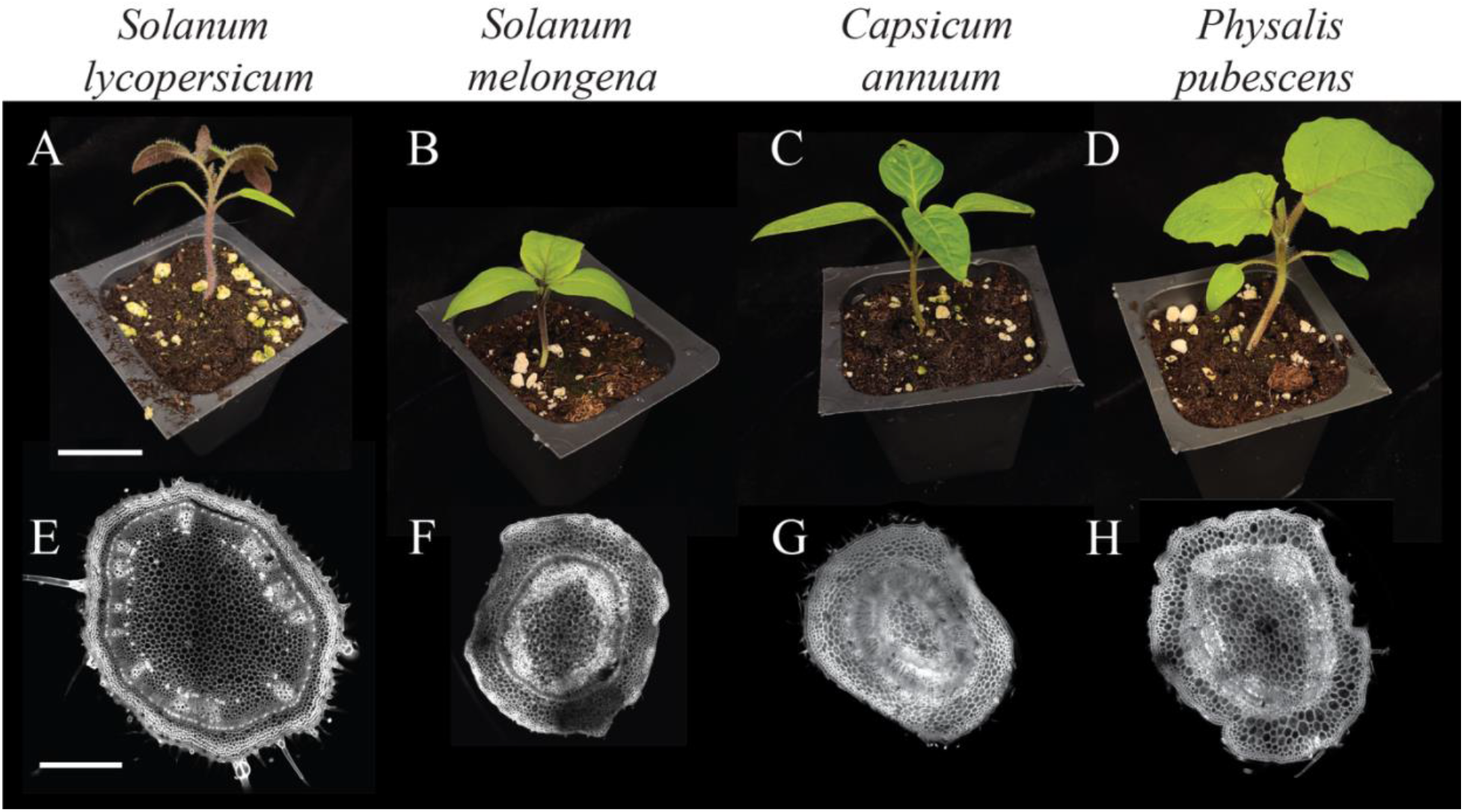
Solanoideae crops share similar stem anatomy. (A-D) Representative images of seedlings at the time of grafting. (E-H) Confocal micrographs show transverse stem sections taken from the location of grafting. *Solanum lycopersicum* (tomato; A, E), *S. melongena* (eggplant; B, F) *Capsicum annuum* (pepper; C, G), and *Physalis pubescens* (groundcherry; D, H) are all members of the Solanoideae sub-family. E-H were stained with propidium iodide and cleared in methyl salicylate prior to confocal imaging. Replication can be found in Figure S1. A-D scale bar = 2 cm, E-H scale bar =1 mm.

We analyzed the overall appearance of the graft combinations 30 days after grafting and observed variable levels of health depending on the species combination. Self-grafts tended to look the healthiest (i.e., turgid and green) (Figures 3A, F, K, P), while groundcherry heterografts generally appeared the least healthy (Figure 3D, H, L M, N, O). Despite this, at least 1 graft survived for each of the 16 combinations, superficially supporting the concept that broad graft compatibility amongst these four species is possible (Figure 4A). Tomato, pepper, and groundcherry self-grafts survived at a high rate (75-95%), while eggplant self-grafts survived only 53% of the time (Figure 4). Notably, this lower survival rate for self-grafted eggplant is consistent with previous work on eggplant grafting (Johnson and Miles, 2011). We observed that most heterograft combinations exhibited moderate rates of survival (40-60%), while eggplant and tomato heterografts had high survival rates (eggplant:tomato (87%), eggplant:pepper (80%)), and multiple combinations with ground cherry as a graft partner showed relatively low survival rates (pepper:groundcherry (20%), eggplant:groundcherry (20%), and groundcherry:eggplant (7%; Figure 4). Curiously, eggplant:pepper was one of the best-surviving heterografts (80%), while the reciprocal pepper:eggplant combination was one of the worst (27%), highlighting the role that directionality might play in determining graft compatibility (Figure 4).

**Figure 3:**
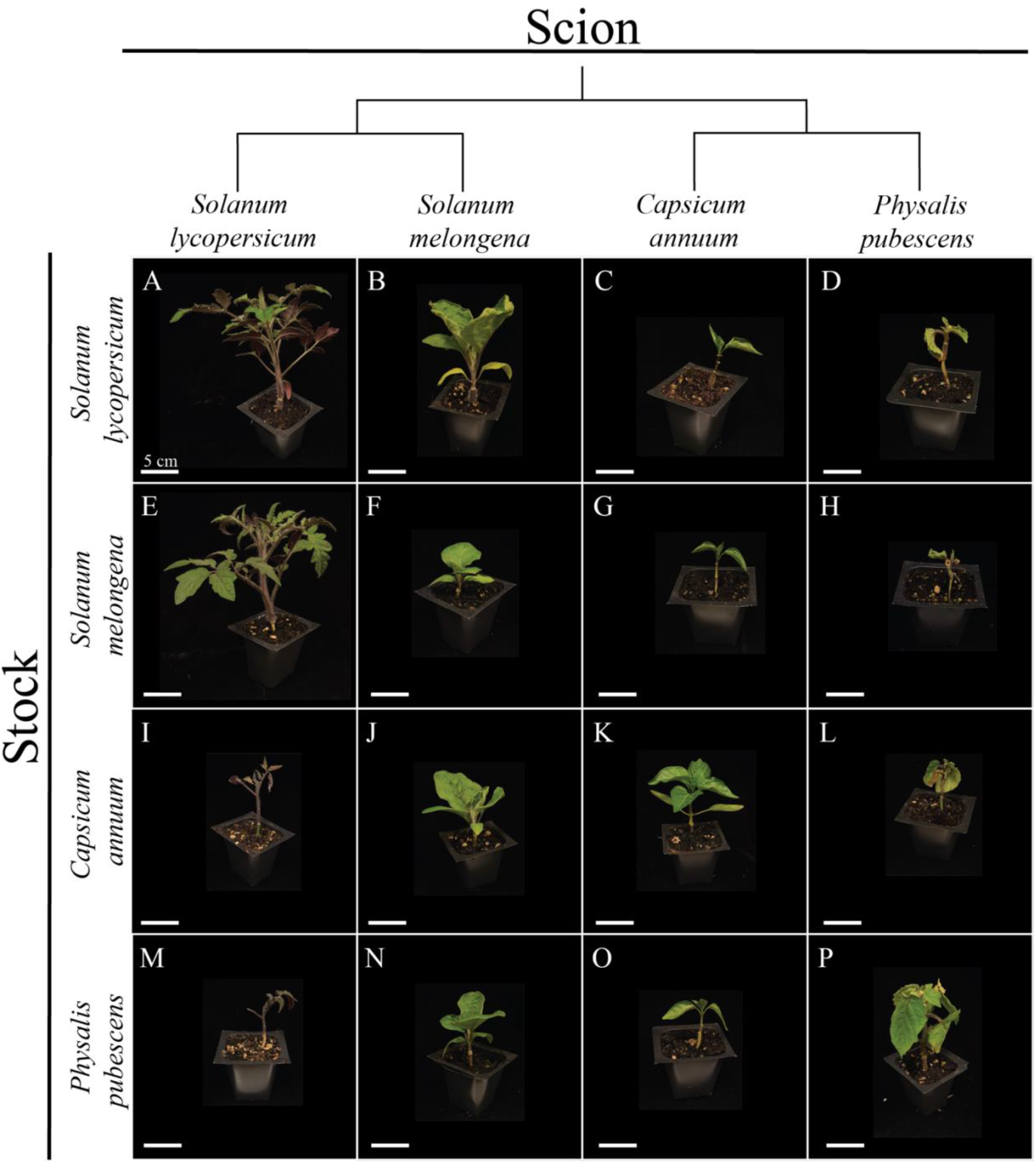
Graft combinations across 4 Solanoideae species 30 DAG. (A-P) Representative images of each graft combination taken at 30 DAG. Tomato:tomato (A), eggplant:tomato (B), pepper:tomato (C), groundcherry:tomato (D), tomato:eggplant (E), eggplant:eggplant (F), pepper:eggplant (G), groundcherry:eggplant (H), tomato:pepper (I), eggplant:pepper (J), pepper:pepper (K), groundcherry:pepper (L), tomato:groundcherry (M), eggplant:groundcherry (N), pepper:groundcherry (O), groundcherry:groundcherry (P). The y-axis of the matrix displays the four stocks: S*olanum lycopersicum* (tomato), *S. melongena* (eggplant), *Capsicum annuum* (pepper), and *Physalis pubescence* (groundcherry). The x-axis also shows the scion species. The phylogenetic relationships of the species are drawn above the scions (Särkinen et al., 2013). Additional replication can be seen in Figure S2. All scale bars = 5 cm.

**Figure 4:**
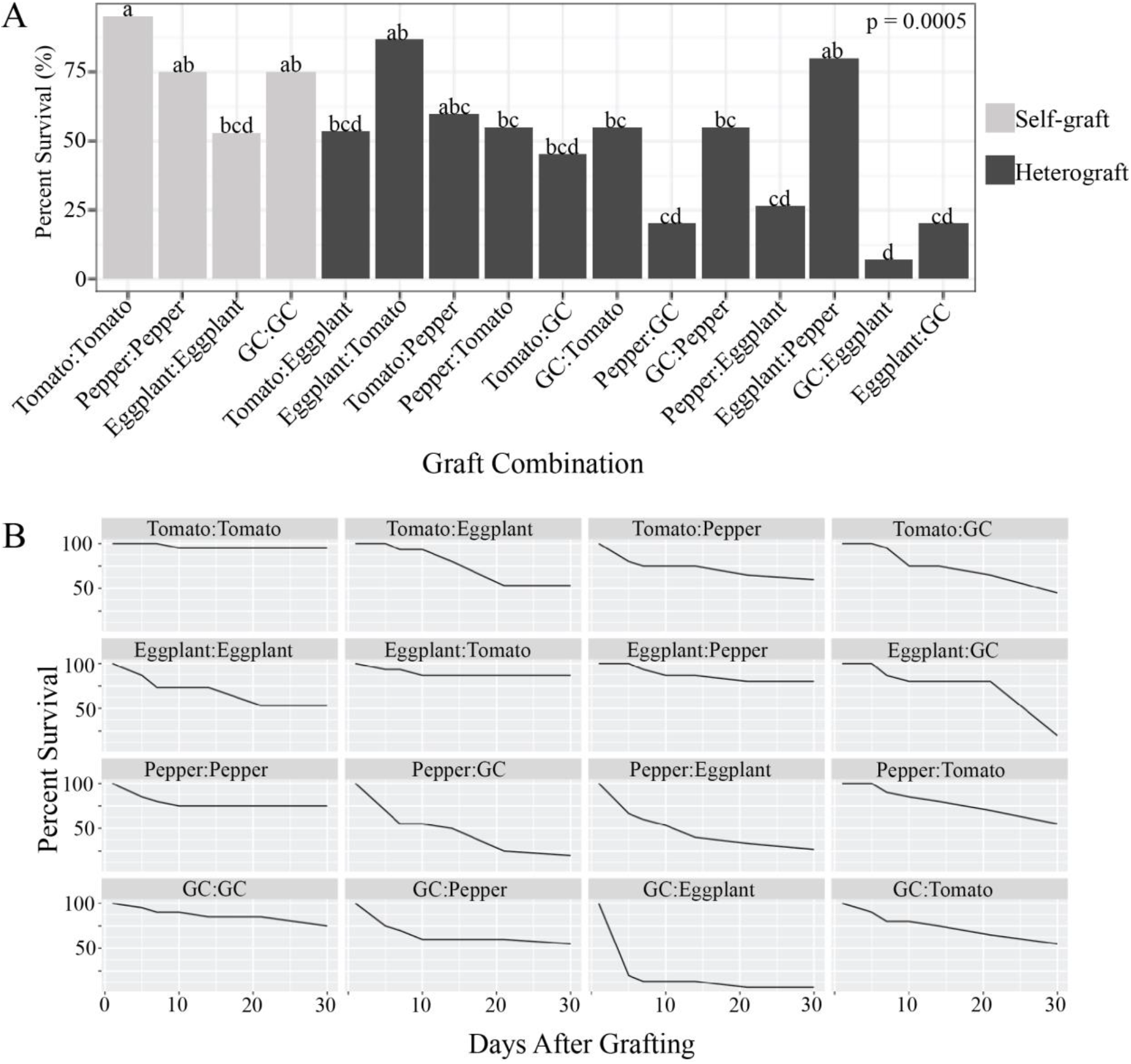
Survival rates for grafted plants 30 DAG. (A) Survival rates 30 DAG for all graft combinations. Light grey bars indicate self-graft combinations, and charcoal bars indicate heterograft combinations. GC = groundcherry. N = 20 for all combinations except those containing eggplant, where n = 15. Compact letters above each bar indicate significant differences between the graft combinations based on pairwise comparisons using Fisher’s Exact Test, *p* < 0.05. (B) Survival rates overtime for all graft combinations. Survival rates were calculated for each graft combination at 5, 7, 10, 14, 21, and 30 DAG. All graft combinations included at least 15 graft replicates; for a detailed description of bioreplication see the materials and methods section.

### 3.2 All intergeneric grafts fail to form vascular reconnections and display delayed incompatibility

To test the qualitative biophysical integrity of the junctions formed in each graft combination, we performed bend tests (Figure 5). The bend test is a simple field test that can be used to measure the integrity of the junction as a proxy for vascular reconnection (Thomas et al., 2022). A well-formed junction is considered to be the strongest part of the stem, as it is packed with lignified vasculature. If a bent stem snaps at the graft site, that is an indicator of poor vascular connectivity, and that plant fails the bend test. If the stem snaps anywhere outside of the junction, the plant passes the bend test. We were unable to break any self-grafted tomato or self-grafted groundcherry junctions, while self-grafted pepper and eggplant junctions broke 5% and 20% of the time, respectively. Most heterografted junctions were easily broken (>94%) except for tomato:eggplant and eggplant:tomato which behaved similarly to self-grafts, breaking at the junction 0% of the time (Figure 5).

**Figure 5:**
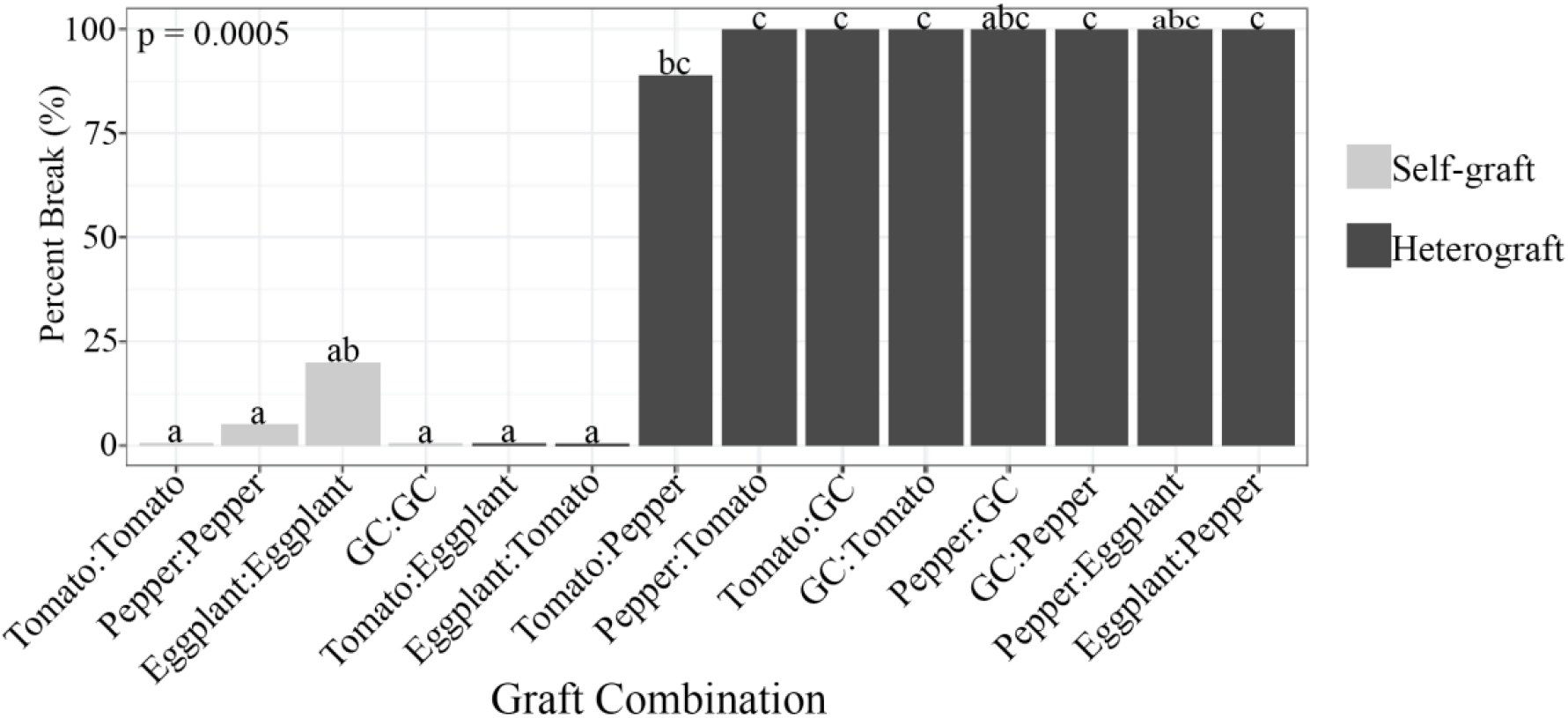
Tomato/eggplant heterografts exhibit strong graft compatibility based on the bend test. The percent of plants that broke at the graft junction during the bend test. Light grey bars indicate self-grafted plants, and dark gray bars indicate heterografted plants. GC = groundcherry. GC:eggplant and eggplant:GC exhibited insufficient survival to conduct the bend test. Compact letters above each bar indicate significant differences between graft combinations based on pairwise comparisons using Fisher’s Exact Test, *p* < 0.05. Replicate values are included in the methods.

To examine the underlying anatomical basis for graft combinations that failed the bend test, we harvested graft junctions from all of our combinations and used confocal microscopy to test for vascular connectivity. Again, groundcherry:eggplant grafts exhibited insufficient survival to collect replicates for image analysis. Our confocal micrographs of self-grafted tomato (Figure 6A, E), eggplant (Figure 6J, M), pepper (Figure 6Q, U), and groundcherry (Figure 6Z, AD) show distinct xylem bridges spanning the graft junction 30 DAG, providing a clear indication of vascular reconnection. Likewise, our tomato:eggplant (Figure 6I, L) and eggplant:tomato (Figure 6B, F) micrographs show well-formed vascular reconnections. In contrast, the other nine graft combinations that we imaged failed to form vascular connections (Figure 6). This result is congruent with our bend tests that indicated graft incompatibility for all of the intergeneric combinations that we tested. Although we could not quantify vascular connectivity for eggplant:groundcherry grafts, the low survival rate for this combination clearly indicates that it is incompatible (Figures 4-5). Thus, we have identified a collection of 9 graft combinations that each exhibit varying survival, with definitive delayed graft incompatibility (Figures 5-6).

**Figure 6:**
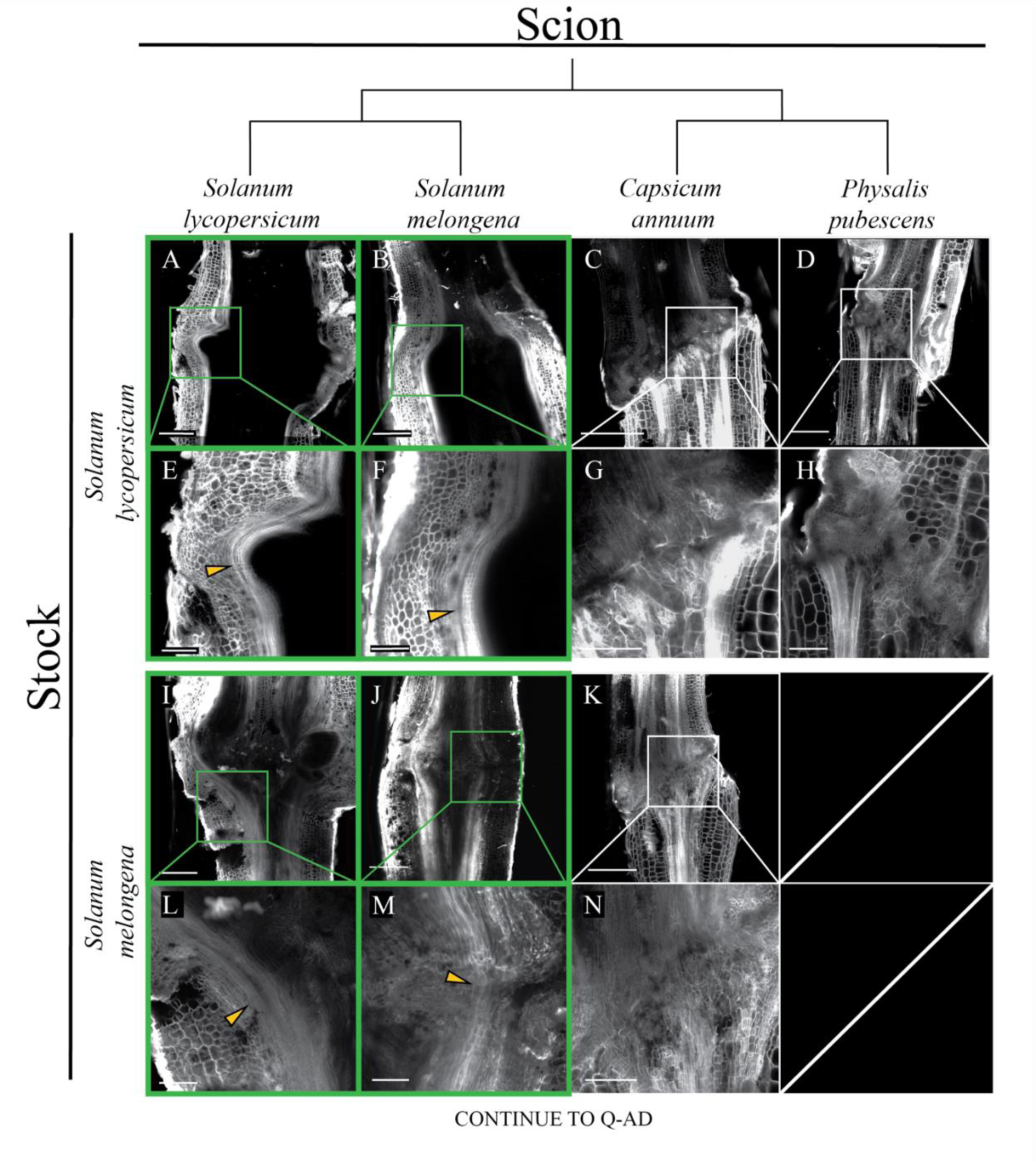
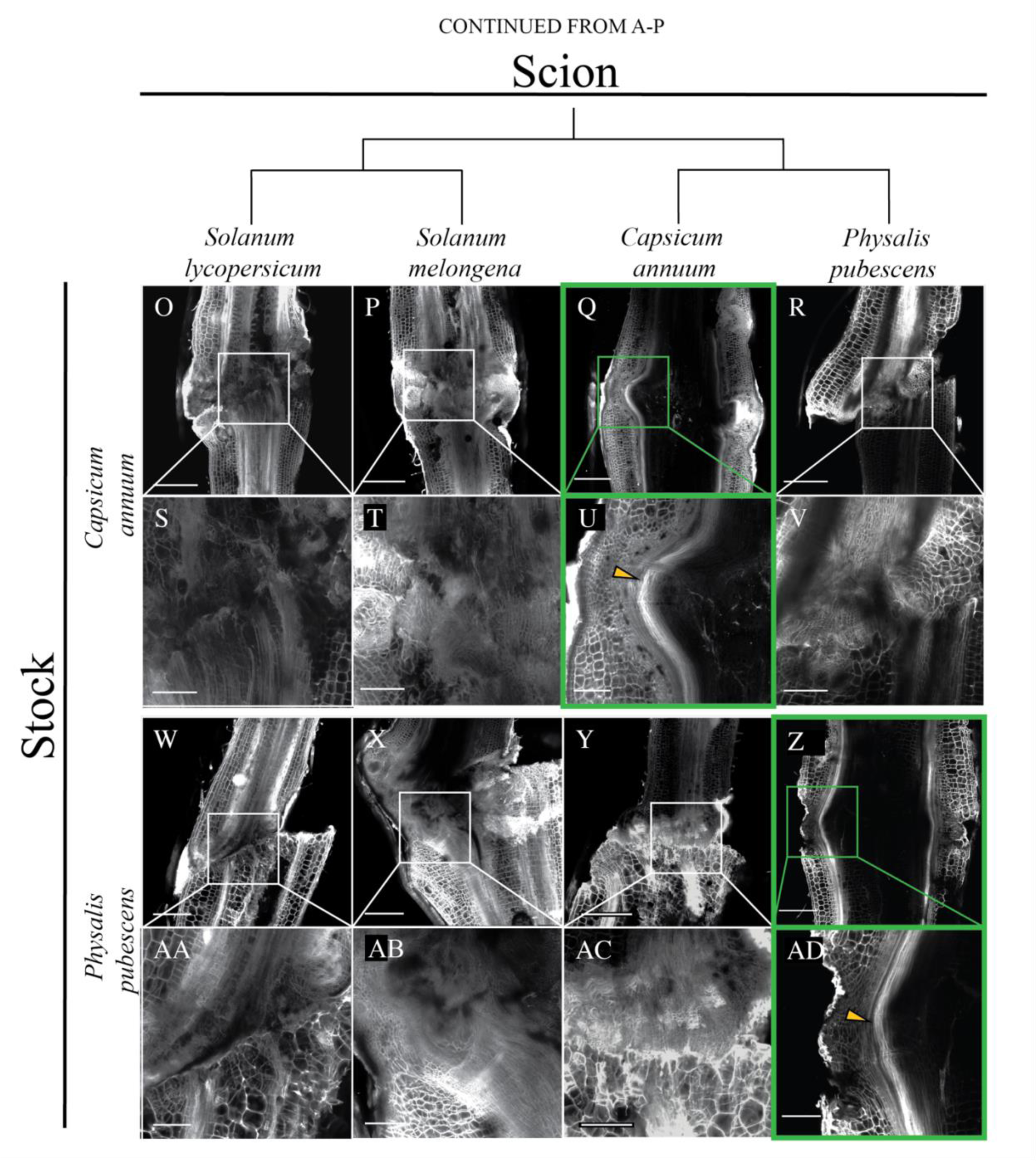
Graft compatibility based on anatomical vascular reconnection is restricted to intrageneric combinations. Confocal micrographs of all surviving graft combinations at 30 DAG. Full junctions for each combination are shown above 3X magnified panels that provide anatomical detail. Tomato:tomato (A,E), eggplant:tomato (B,F), pepper:tomato (C,G), groundcherry:tomato (D,H), tomato:eggplant (I,M), eggplant:eggplant (J,N), pepper:eggplant (K,O), tomato:pepper (O,S), eggplant:pepper (P,T), pepper:pepper (Q,U), groundcherry:pepper (R,V), tomato:groundcherry (W,AA), eggplant:grounchcherry (X,AB) pepper:groundcherry (Y,AC), groundcherry:groundcherry (Z,AD). Groundcherry:eggplant grafts exhibited insufficient survival to be imaged with statistically relevant replication. The y-axis of the image matrix displays the four stocks: S*olanum lycopersicum* (tomato), *S. melongena* (eggplant), *Capsicum annuum* (pepper), and *Physalis pubescence* (groundcherry). The x-axis shows the scion genotypes. The phylogenetic relationship of the scions is shown along the x-axis above the scions (Särkinen et al., 2013). Panels outlined with green boxes indicate grafts with successful vascular bridges, and panels outlined with white boxes indicate grafts with failed vascular connections. Yellow arrows point to xylem bridges. Grafts were harvested, stained with propidium iodide, and cleared in methyl salicylate prior to imaging. A-D, I-K, O-R, and W-Z scale bars are 1 mm. E-H, L-N, S-V, and AA-AD scale bars are 333 μm. Additional replicates are included in Figure S3.

To investigate whether the incompatible grafts in our study exhibit other measurable symptoms of graft failure, we measured scion and stock growth rates. Our data shows significant differences in lateral growth between all graft combinations based on changes in stem diameter at 30 DAG (Scion: F_15,155_=30.68, p<2e-16; Stock:F_15,155_=27.27, p<2e-16; Figure 7A-B, Table S1-3). We first wanted to examine the relationship between self-and heter1ografted stems. Using an ANOVA, we showed that there are significant differences in lateral growth between self-and heterografted stems, with self-grafts generally exhibiting healthier growth, where scions were 148% wider (Scion: F_1,155_=51.83, p<2.51e-11), and stocks were 91% wider than their heterografted counterparts (F_1,155_=25.64, p<1.16e-06; Figure 7A-B, Table S1-4).

**Figure 7:**
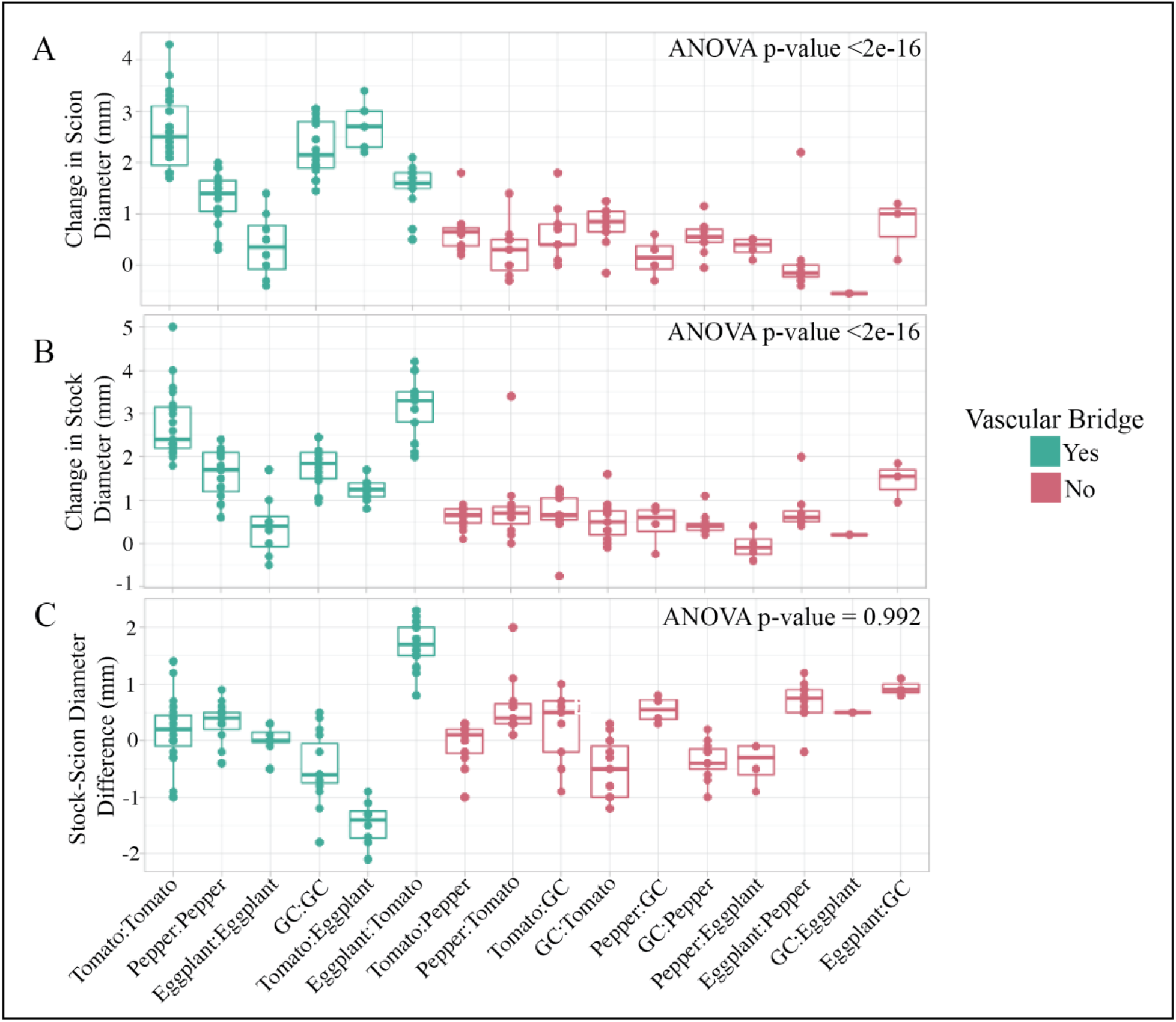
Stem growth is correlated with graft compatibility. (A) The change in scion diameter 30 DAG for all graft combinations. (B) The change in stock diameter 30 DAG for all graft combinations. (C) The difference between stock and scion diameter 30 DAG for all graft combinations. This differential growth was calculated by subtracting the scion diameter from the stock diameter of each graft combinations 30 DAG. Teal indicates compatible grafts that formed successful vascular bridges (Figure 6), red indicates incompatible grafts that failed to form vascular reconnections. ANOVA was calculated between compatible and incompatible plants, p-value <0.05.

Since our anatomical and biophysical data indicate that eggplant:tomato and tomato:eggplant heterografts are compatible, we next tested if graft compatibility is more strongly correlated with increased lateral stem and stock growth post-grafting. We found a significant, pronounced difference in stem diameter, wherein compatible scions and stock were 392% (F_1,155_=139.7, p<2e-16) and 213% (F_1,155_=107, p<2e-16) wider than their incompatible counterparts, respectively (Figure 7A-B). Therefore the most significant variance that we detected in lateral growth was correlated with graft compatibility.

In woody crops, bulging scions are commonly noted as symptoms of graft incompatibility (Andrews and Marquez, 2010). To test differential growth between the scion and stock of individuals plants, we looked at the diameter of the stock minus the diameter of the scion 30 DAG. While many of the heterografted combinations had varying diameters between the scion and stock (Figure 7C), we found no significant relationship between junction bulging and graft incompatibility (F_1,155_=0, p = 0.992; Table S4).

In conclusion, we found that while Solanoideae plants have a broad ability to form non-vascular callus connections, very few of the plants are truly compatible (Figure 6). We show that the bend test is a reliable, fast, and low-tech test for graft compatibility (Figure 5). Tomato and eggplant heterografts survive and build vascular connections similarly, if not better, to their self-grafted counterparts, making this combination an excellent choice for hetero-compatible graft studies (Figures 2-6). We also demonstrate that comparing lateral growth of herbaceous stems can act as an early predictor for graft incompatibility (Figure 7). Furthermore, we highlight the importance of graft-compatible studies, as survival is not a reliable indicator of true graft success.

## 4. DISCUSSION

The Solanaceae family is often noted as a highly graftable group of plants with eggplant, potato, and tobacco all capable of grafting with tomato (Lee and Oda, 2010; Dawson, 1942; Notaguchi et al., 2020b). To explore this statement, we conducted an in-depth analysis of reciprocal grafts using four agronomically relevant crops: tomato (*Solanum lycopersicum*), pepper (*Capsicum annuum*), eggplant (*S. melongena*), and groundcherry (*Physalis pubescens*; Figs 1-2). All four of these species belong to the same sub-family, making these four crops more closely related than previous studies looking at graft compatibility between Nicotianideae and Solanoideae (Notaguchi et al., 2015, 2020b).

We performed reciprocal grafts amongst all four species, leading to the production of 16 graft combinations (Figure 3). We observed a variety of survival rates at 30 DAG; self-grafted plants had high survival rates, except for eggplant which performed as previously predicted (Figure 4; Johnson and Miles, 2011), while most heterografts survived at moderate rates (40-60% survival; Figure 4). Interestingly, one of the highest surviving heterografted plants 30 DAG was eggplant:pepper. Further investigation into this graft showed that despite eggplant:pepper surviving in high numbers (80%), no vascular bridges were present in the graft junction, phenocopying the graft junction of many other incompatible graft combinations in Solanaceae such as tomato-pepper grafts (Figure 6; Thomas et al., 2022).

The ability of non-vascular tissue to heal and sustain life, in the absence of vascular reconnection is the definition of delayed incompatibility (Argles, 1937). The fact that plants without continuous vascular strands are able to survive, raises a whole suite of interesting questions regarding the physiological requirements for sustained life.

All 16 graft combinations produced at least 1 surviving graft, while only six produced vascular bridges (Figures 4-6). Out of the six compatible grafts, four were self-grafts, and only tomato:eggplant and eggplant:tomato were compatible heterografts (Figure 6). These findings demonstrate the importance of looking beyond graft survival to determine whether a given combination is truly compatible. Further testing, including biostability or anatomical analysis is required for confident characterization of graft compatibility. Bend tests further bolstered our findings that although many of the graft combinations survived at rates similar to compatible grafts, the integrity of the stem was compromised due to failed vascular reconnection.

By selecting species from different genera, we were able to identify compatibility constraints within the Solanoideae sub-family. As we expected, we found that all self-grafts were compatible. However, the intrageneric grafts that we performed between tomato and eggplant were the only heterografts with true compatibility. While these two species did diverge around 14 ma, they are both members of the *Solanum* genus. In addition, potato (*Solanum tuberosum*) is also compatible with tomato and eggplant, further supporting intrageneric compatibility within *Solanum* (Thompson and Morgan, n.d.). All of the other intergeneric grafts that we tested, regardless of survival, were incompatible. Previous work investigating Solanaceae rootstocks that could be used for eggplant scion production, showed that species outside of *Solanum* are incompatible with eggplant. We were able to extend this model for incompatibility, by identifying intergeneric graft limitations between *Solanum, Capsicum*, and *Physalis (Ali et al. 1990)*.

While further graft studies are required to definitively state that intergeneric grafts are not possible within Solanoideae, our work demonstrates that phylogenetic constraints play a role in determining graft compatibility. The mechanisms underlying phylogenetic limitations to grafting have yet to be determined; however, we hypothesize that failed communication is responsible for this observed incompatibility. Future studies looking into the role of intercellular signals, as well as innate immune responses will help elucidate the precise mechanisms that define the evolutionary boundaries of graft compatibility, and inform the predictive selection and expansion of successful graft partners.

## 5. CONCLUSIONS

In this study, we show that intergeneric grafting with four species from the Solanoideae produces multiple instances of delayed incompatibility. Furthermore, we demonstrate that heterografts between tomato and eggplant form truly compatible junctions, indicated by vascular reconnections that form biophysically stable grafts. Together, these compatible and incompatible graft combinations provide a useful toolkit to explore the underlying genetic mechanisms for graft compatibility. Our research provides strong support showing that graft compatibility within the Solanoideae is limited to intrageneric species combinations. Future work using additional Solanaceous species can be used to test whether intrageneric graft limitations exist broadly across the family.

## AUTHOR CONTRIBUTIONS

H.R.T and M.H.F. conceived and designed the study. H.R.T. and A.G.. gathered experimental data. H.R.T. analyzed the data. H.R.T., A.G., and M.H.F. wrote the manuscript.

## ACKNOWLEDGMENTS

The authors thank the Cornell Growth Chamber Facility and the Cornell Institute of Biotechnology Imaging Facility for their assistance. This work was supported by Frank Lab startup funds from Cornell University College of Agriculture and Life Sciences. M.H.F was supported by the National Science Foundation (NSF) (CAREER IOS-1942437). H.R.T. was supported by a United States Department of Agriculture National institute of Food and Agriculture (USDA-NIFA) Predoctoral Fellowship (2020-67011-31882); A.G. utilized funding from the Cornell Institute for Digital Agriculture Research Innovation Fund. Image data was acquired through the Cornell Institute of Biotechnology’s Imaging Facility, with NIH (S10OD018516) funding for the shared Zeiss LSM880 confocal/multiphoton microscope.

## SUPPORTING INFORMATION

Table S1: Table- Growth of grafted plants 0-30 DAG.

Table S2: Table- Tukey Multiple Comparison of Means of scion diameter 30 DAG..

Table S3. Table- Tukey Multiple Comparison of Means of stock diameter 30 DAG.

Table S4: Table- ANOVA Results.

Table S5. Table- Levene’s and Wilks-Shapiro Tests for ANOVA assumptions.

Figure S1: Solanoideae species utilized in graft trial.

Figure S2: Solanoideae graft combinations 30 DAG.

Figure S3: Propidium iodide stained solanoideae graft junctions 30 DAG.

Figure S4: Tukey’s Multiple Comparison of Means, Levene’s Test and Wilks-Shapiro Tests.

## REFERENCES

Ali, M., Mohammad, A.L.I., and Fujieda, K. (1990). Cross Compatibility between Eggplant (Solanum melongena L.) and Wild Relatives. Journal of the Japanese Society for Horticultural Science 58: 977–984.

Andrews, P.K. and Marquez, C.S. (2010). Graft Incompatibility. Horticultural Reviews: 183–232.

Argles, G.K. (1937). A Review of the Literature on Stock-scion Incompatibility in Fruit Trees: With Particular Reference to Pome and Stone Fruits.

Benda, G.T.A., Hartmann, H.T., and Kester, D.E. (1960). Plant Propagation: Principles and Practices. American Midland Naturalist 63: 253.

Dawson, R.F. (1942). ACCUMULATION OF NICOTINE IN RECIPROCAL GRAFTS OF TOMATO AND TOBACCO. American Journal of Botany 29: 66–71.

Errea, P., Felipe, A., and Herrero, M. (1994). Graft establishment between compatible and incompatiblePrunusspp. Journal of Experimental Botany 45: 393–401.

Fox, J. and Weisberg, S. (2018). An R Companion to Applied Regression (SAGE Publications).

Frodin, D.G. (2004). History and concepts of big plant genera. TAXON 53: 753–776.

Goldschmidt, E.E. (2014). Plant grafting: new mechanisms, evolutionary implications. Frontiers in Plant Science 5.

Hériché, M., Arnould, C., Wipf, D., & Courty, P. E. (2022). Imaging plant tissues: advances and promising clearing practices. Trends in Plant Science.

Hervé, M. (2020) Package ‘RVAideMemoire. https://CRANR-projectorg/package=RVAideMemoire.

Johnson, S.J. and Miles, C.A. (2011). Effect of Healing Chamber Design on the Survival of Grafted Eggplant, Tomato, and Watermelon. HortTechnology 21: 752–758.

Kawaguchi, M., Taji, A., Backhouse, D., and Oda, M. (2008). Anatomy and physiology of graft incompatibility in solanaceous plants. The Journal of Horticultural Science and Biotechnology 83: 581–588.

Kubota, C., McClure, M.A., Kokalis-Burelle, N., Bausher, M.G., and Rosskopf, E.N. (2008). Vegetable Grafting: History, Use, and Current Technology Status in North America. HortScience 43: 1664–1669.

Lee, J.-M. and Oda, M. (2010). Grafting of Herbaceous Vegetable and Ornamental Crops. Horticultural Reviews: 61–124.

Liu, N., Zhou, B., Zhao, X., Lu, B., Li, Y., and Hao, J. (2009). Grafting Eggplant onto Tomato Rootstock to Suppress Verticillium dahliae Infection: The Effect of Root Exudates. HortScience 44: 2058–2062.

Melnyk, C.W. (2017). Plant grafting: insights into tissue regeneration. Regeneration 4: 3– 14.

Mudge, K., Janick, J., Scofield, S., and Goldschmidt, E.E. (2009a). A History of Grafting. Horticultural Reviews: 437–493.

Mudge, K., Janick, J., Scofield, S., and Goldschmidt, E.E. (2009b). A History of Grafting. Horticultural Reviews: 437–493.

Notaguchi, M. et al. (2020a). Cell-cell adhesion in plant grafting is facilitated by β-1,4-glucanases. Science 369: 698–702.

Notaguchi, M. et al. (2020b). Cell-cell adhesion in plant grafting is facilitated by β-1,4-glucanases. Science 369: 698–702.

Notaguchi, M., Higashiyama, T., and Suzuki, T. (2015). Identification of mRNAs that move over long distances using an RNA-Seq analysis of Arabidopsis/Nicotiana benthamiana heterografts. Plant Cell Physiol. 56: 311–321.

R Core Team (2021). R: A language and environment for statistical computing. R Foundation for Statistical Computing.

Rasool, A., Mansoor, S., Bhat, K.M., Hassan, G.I., Baba, T.R., Alyemeni, M.N., Alsahli, A.A., El-Serehy, H.A., Paray, B.A., and Ahmad, P. (2020). Mechanisms Underlying Graft Union Formation and Rootstock Scion Interaction in Horticultural Plants. Front. Plant Sci. 11: 590847.

Reeves, G., Tripathi, A., Singh, P., Jones, M.R.W., Nanda, A.K., Musseau, C., Craze, M., Bowden, S., Walker, J.F., Bentley, A.R., Melnyk, C.W., and Hibberd, J.M. (2022). Monocotyledonous plants graft at the embryonic root–shoot interface. Nature 602: 280–286.

Romano, D. and Paratore, A. (2001). EFFECTS OF GRAFTING ON TOMATO AND EGGPLANT. Acta Horticulturae: 149–154.

Särkinen, T., Bohs, L., Olmstead, R.G., and Knapp, S. (2013). A phylogenetic framework for evolutionary study of the nightshades (Solanaceae): a dated 1000-tip tree. BMC Evol. Biol. 13: 214.

Stern, S. and Bohs, L. (2012). An explosive innovation: Phylogenetic relationships of Solanum section Gonatotrichum (Solanaceae). PhytoKeys: 89–98.

Thomas, H., Van den Broeck, L., Spurney, R., Sozzani, R., and Frank, M. (2022). Gene regulatory networks for compatible versus incompatible grafts identify a role for SlWOX4 during junction formation. Plant Cell 34: 535–556.

Thompson and Morgan (n.d.) TomTato® (Ketchup ‘n’ Fries™, Ketchup and Chips). Thompson and Morgan.

Westwood, M.N. (1993). Temperate-zone Pomology: Physiology and Culture (Timber Press (OR)).

Zeist, A.R., Giacobbo, C.L., da Silva Neto, G.F., Zeist, R.A., da R Dorneles, K., and de Resende, J.T.V. (2018). Compatibility of tomato cultivar Santa Cruz Kada grafted on different Solanaceae species and control of bacterial wilt. Horticultura Brasileira 36: 377–381.

