## Supplemental Figures for "Anatomical and biophysical characterization of intergeneric graft-incompatibility within the Solanoideae"

**SUPPORTING INFORMATION**

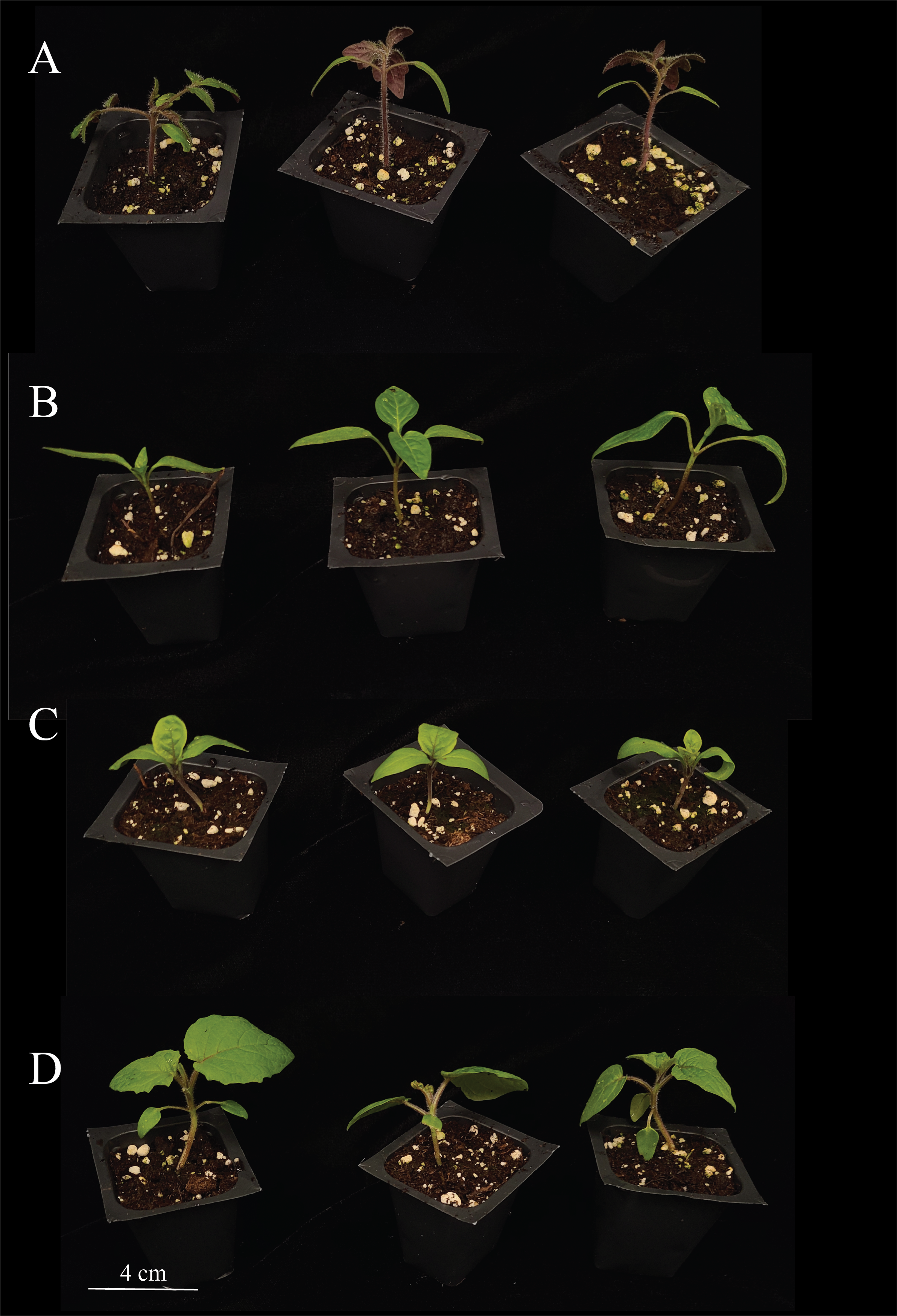

**Figure S1: Solanoideae species utilized in graft trial.** Duplication of tomato (A), pepper (B), eggplant (C), and groundcherry (D) seedlings used in the graft trial (Figures 2). All images set to equal, scale bar = 4 cm.

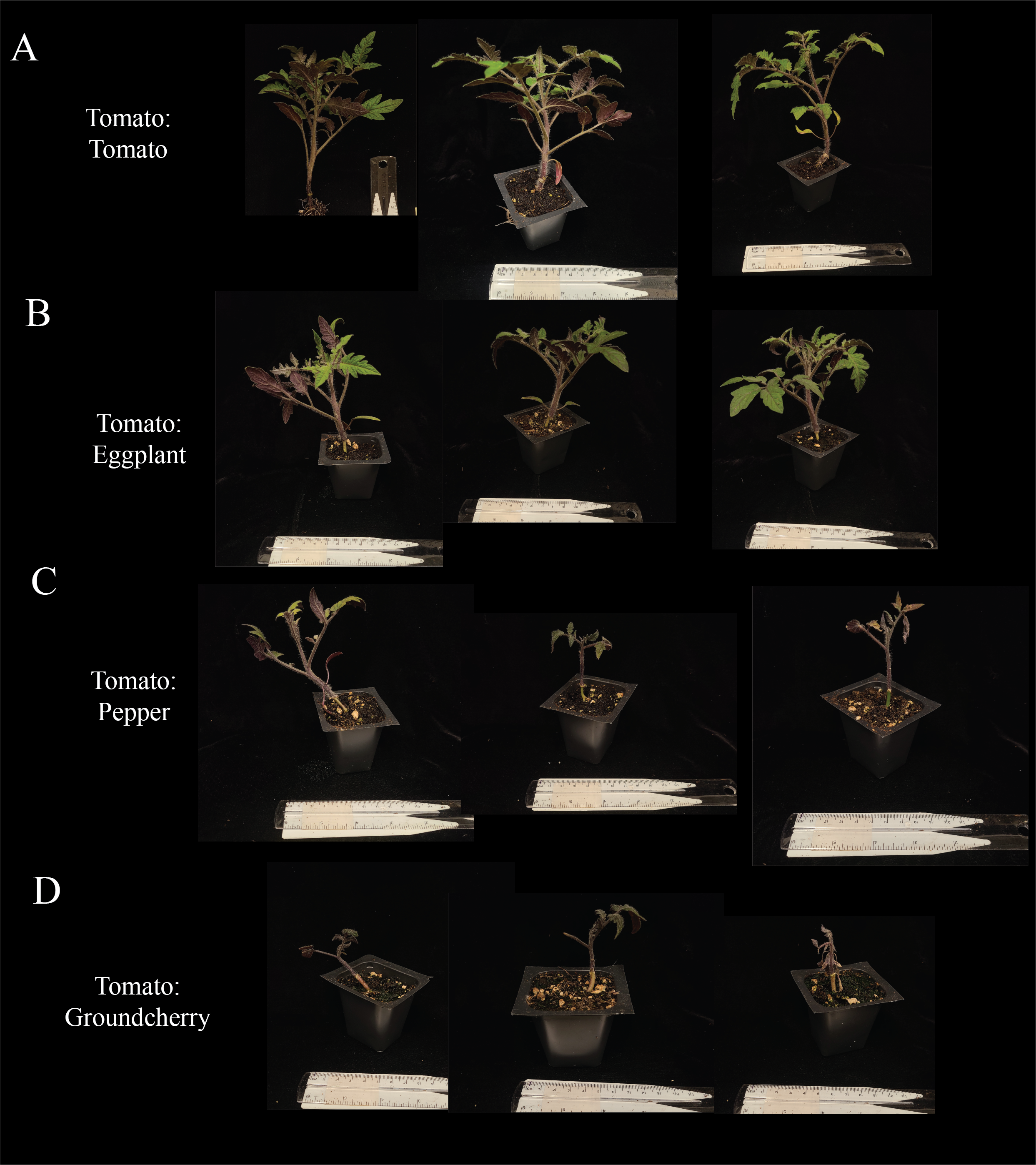

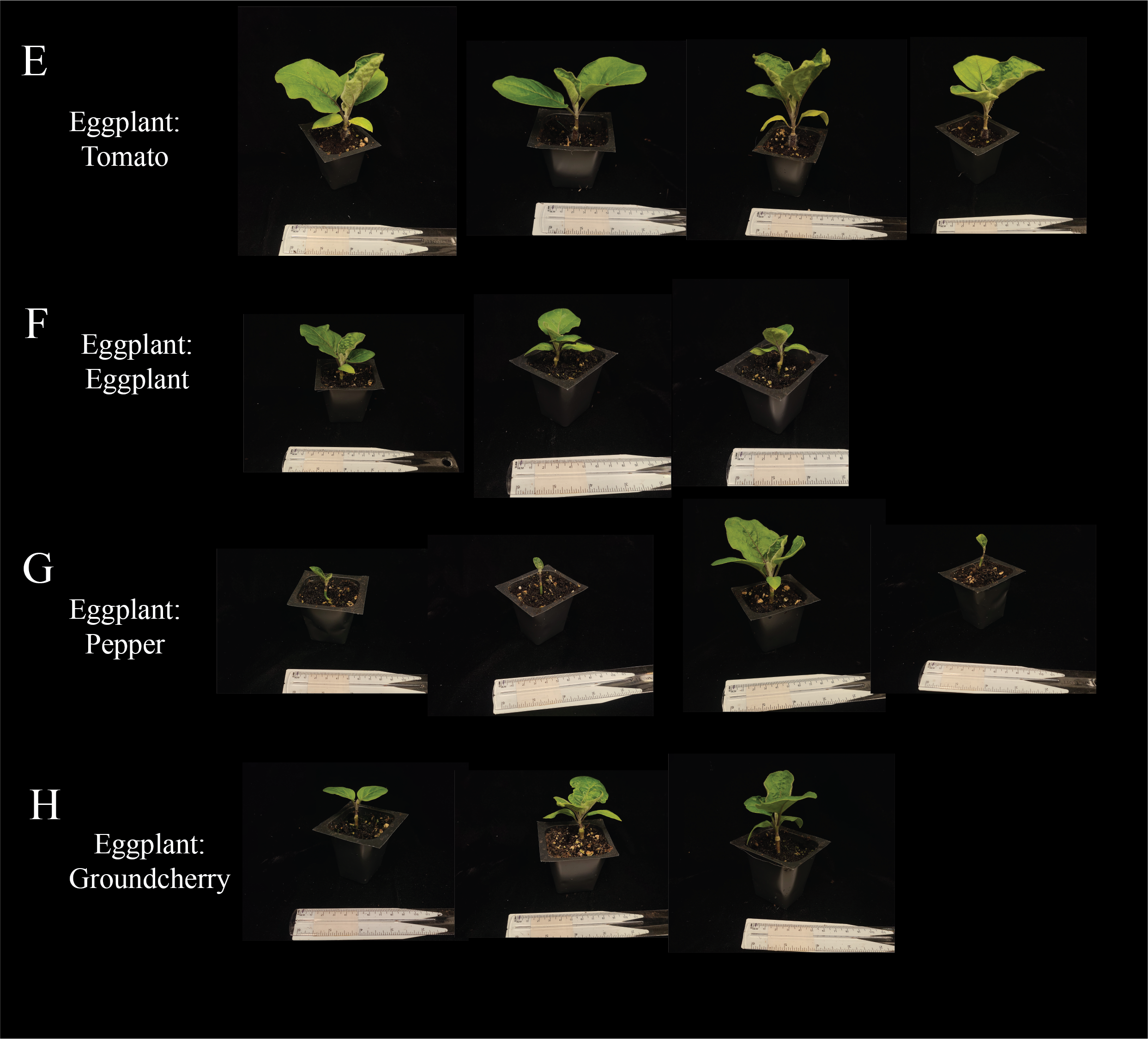

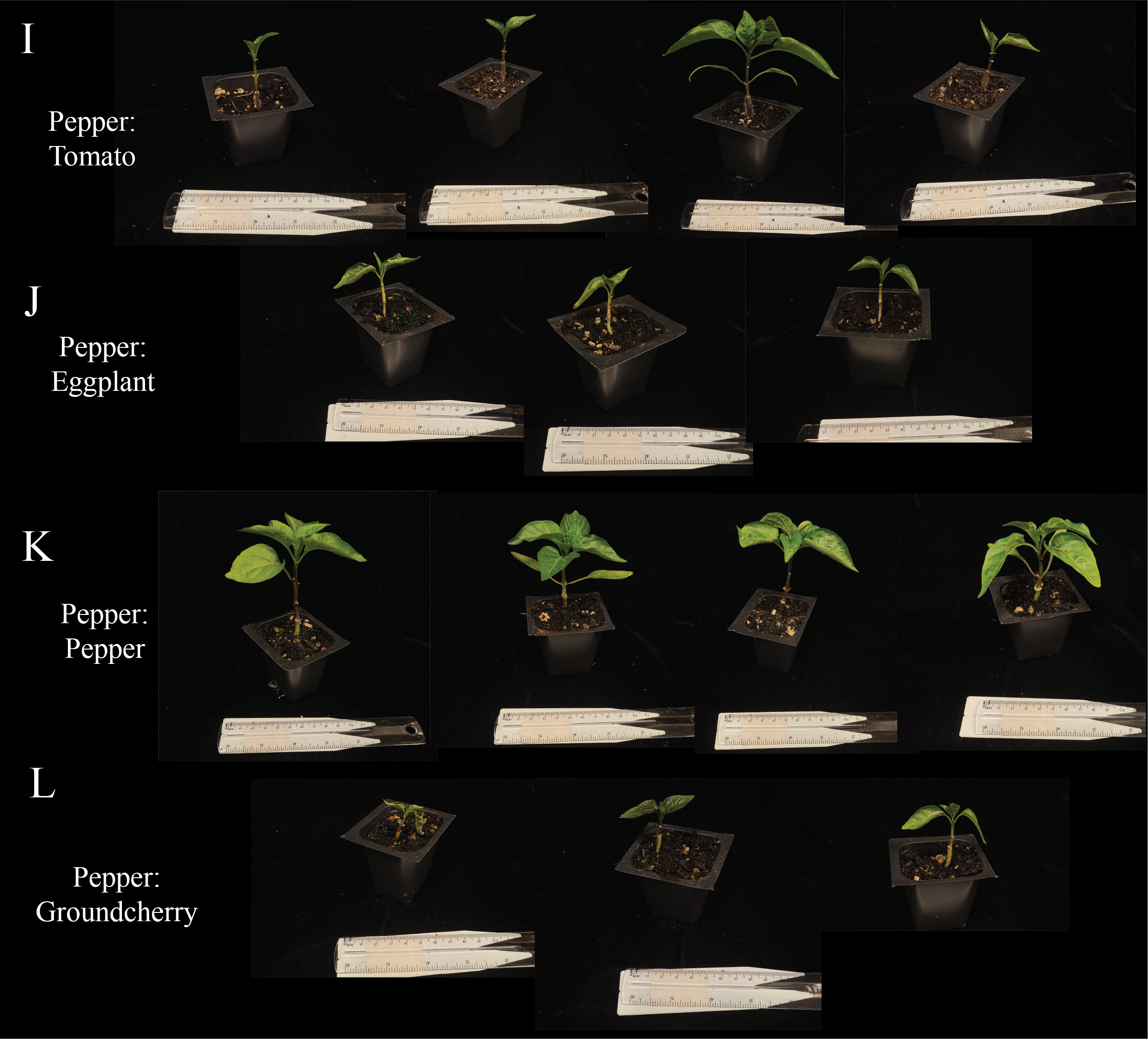

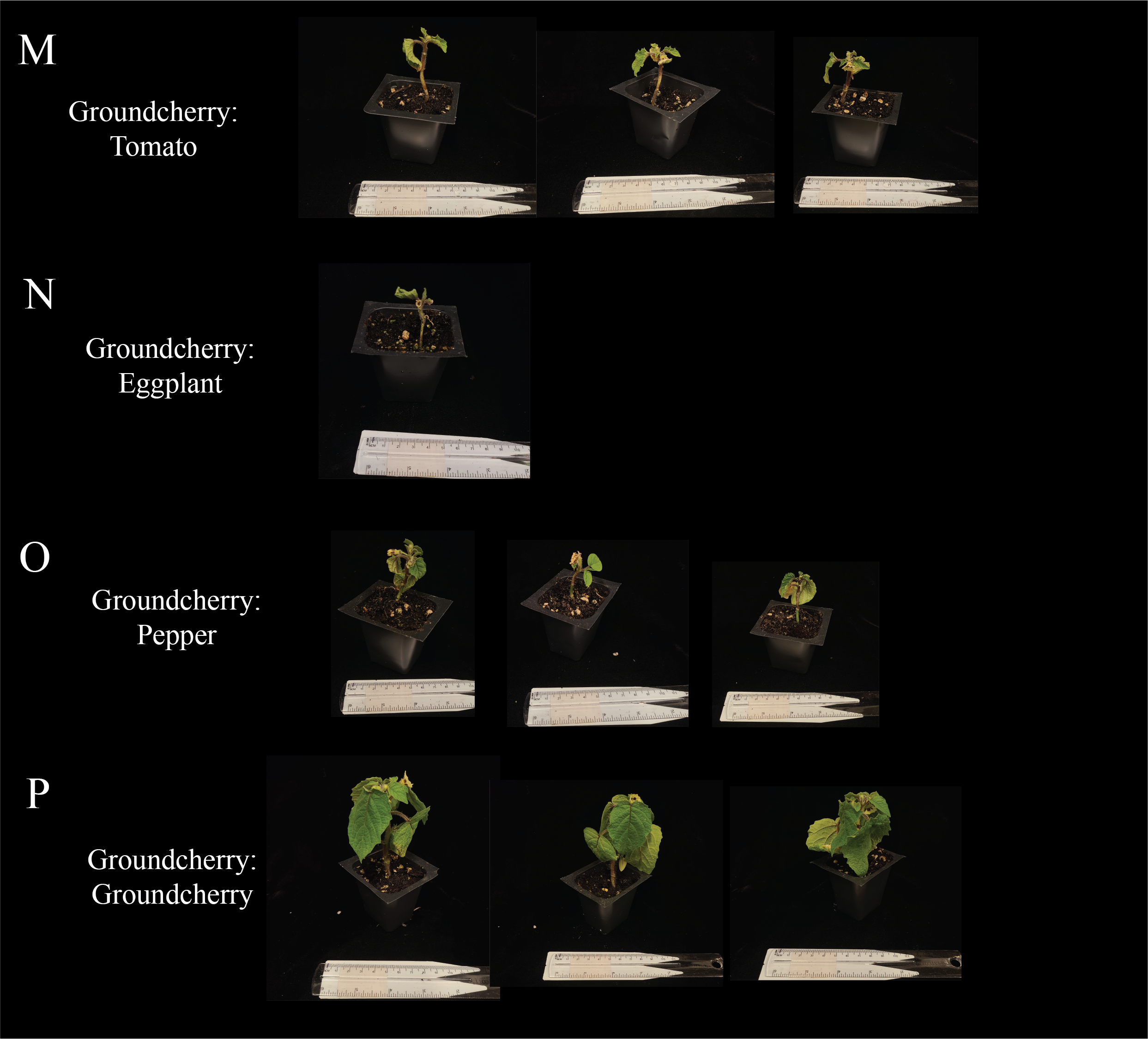

**Figure S2: Solanoideae graft combinations 30 DAG.** Duplication of tomato:tomato (A), tomato:eggplant (B), tomato:pepper (C ), tomato:groundcherry (D), eggplant:tomato (E) , eggplant:eggplant (F), eggplant:pepper (G), eggplant:groundcherry (H), pepper:tomato (I), pepper:eggplant (J), pepper:pepper (K), pepper:groundcherry (L), groundcherry:tomato (M), groundcherry:eggplant (N), groundcherry:pepper (O), groundcherry:groundcherry (P) grafts shown in Figure 3.Ruler shown below each plant.

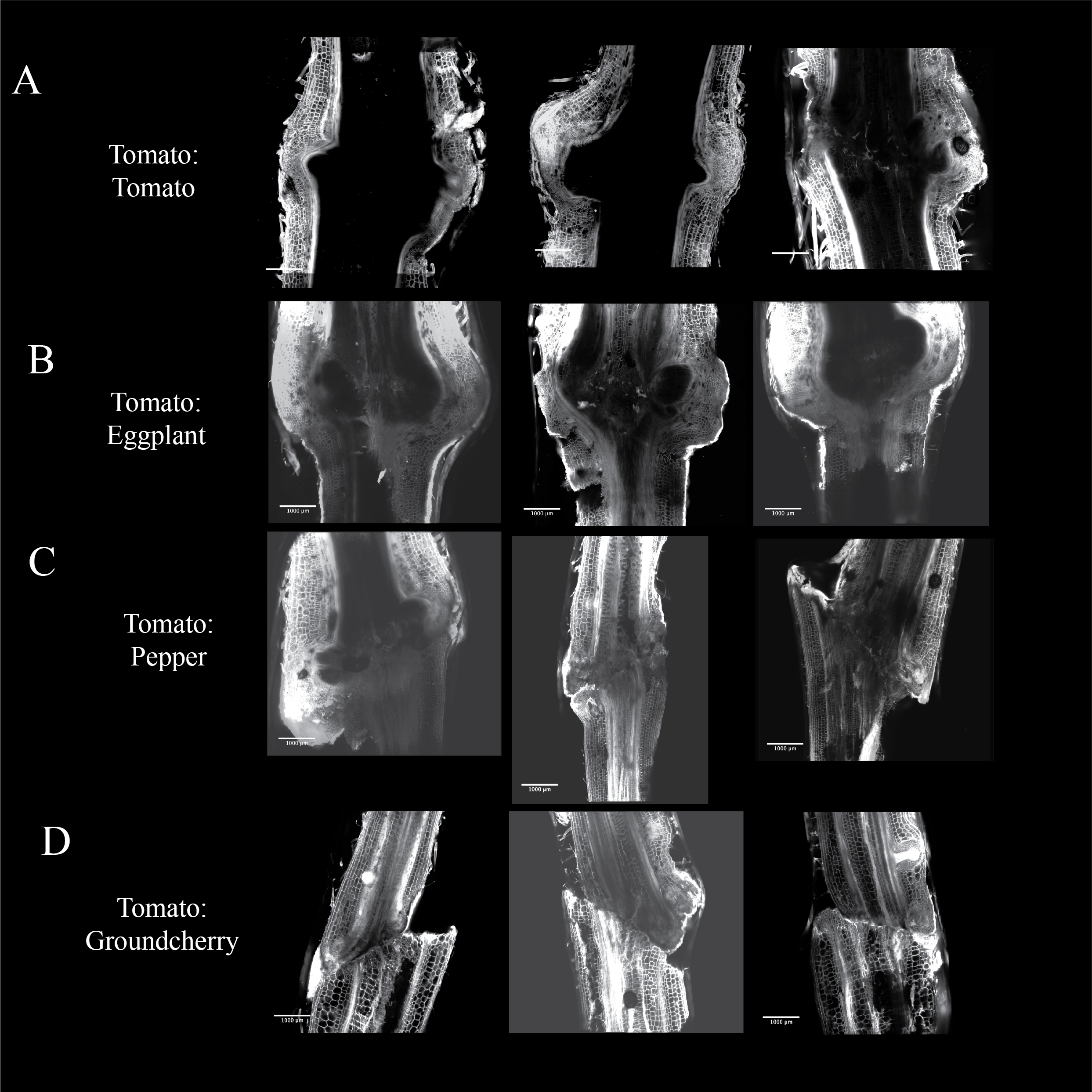

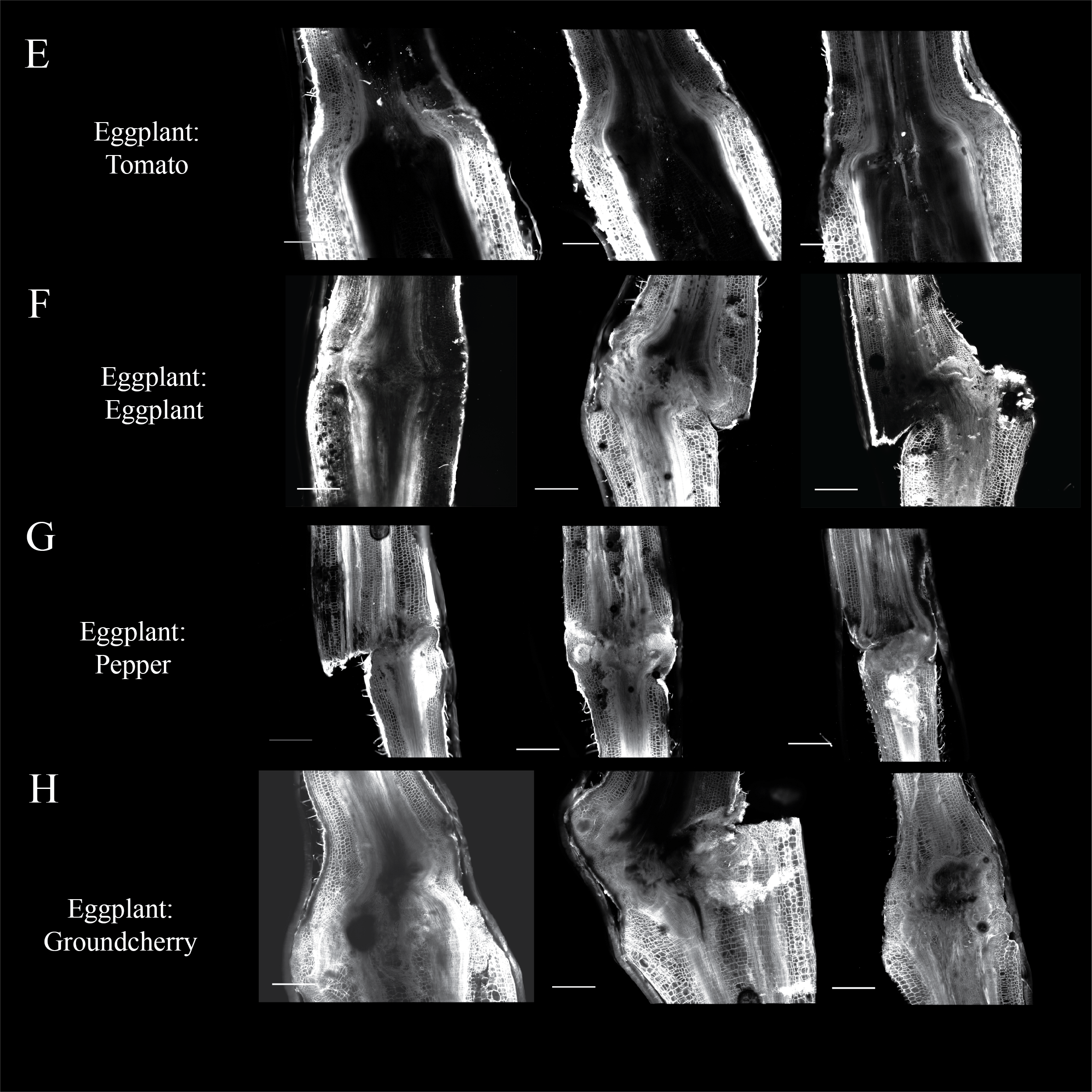

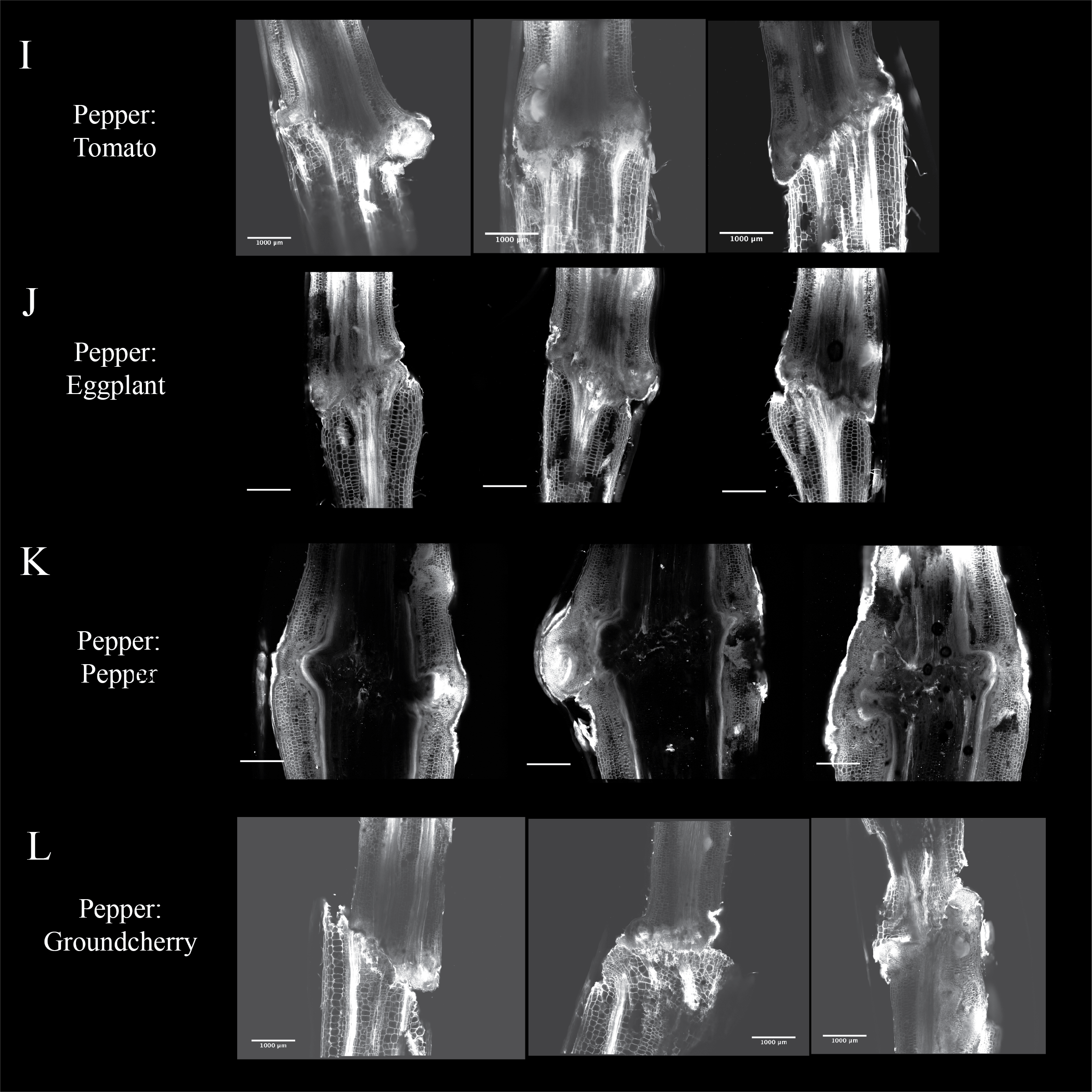

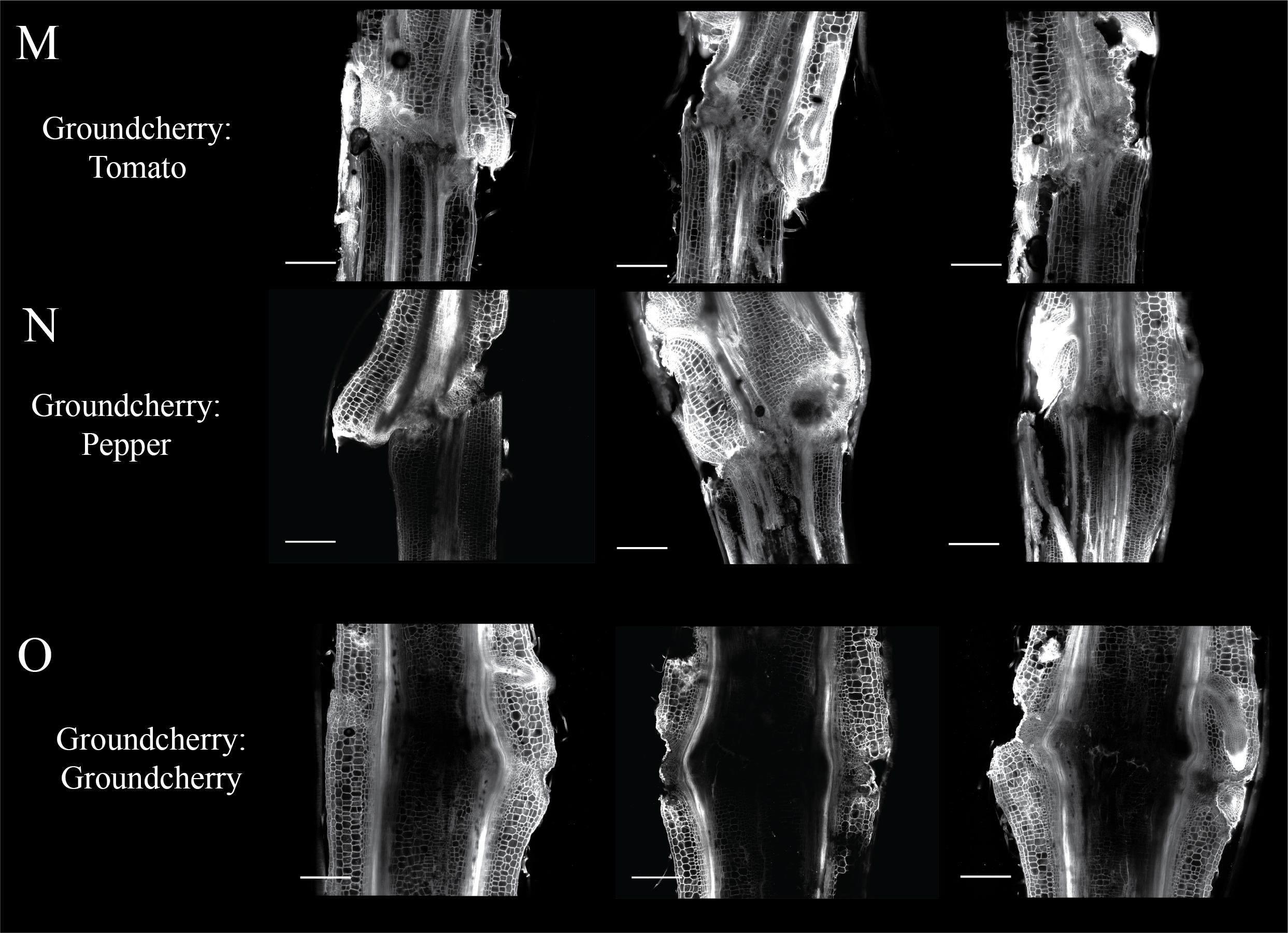

**Figure S3: Propidium iodide stained solanoideae graft junctions 30 DAG.** Duplication of tomato:tomato (A), tomato:eggplant (B), tomato:pepper (C ), tomato:groundcherry (D), eggplant:tomato (E) , eggplant:eggplant (F), eggplant:pepper (G), eggplant:groundcherry (H), pepper:tomato (I), pepper:eggplant (J), pepper:pepper (K), pepper:groundcherry (L), groundcherry:tomato (M), groundcherry:eggplant (N), groundcherry:pepper (O), groundcherry:groundcherry (P) grafts shown in Figure 6. Scale bars are equal to 1 mm.

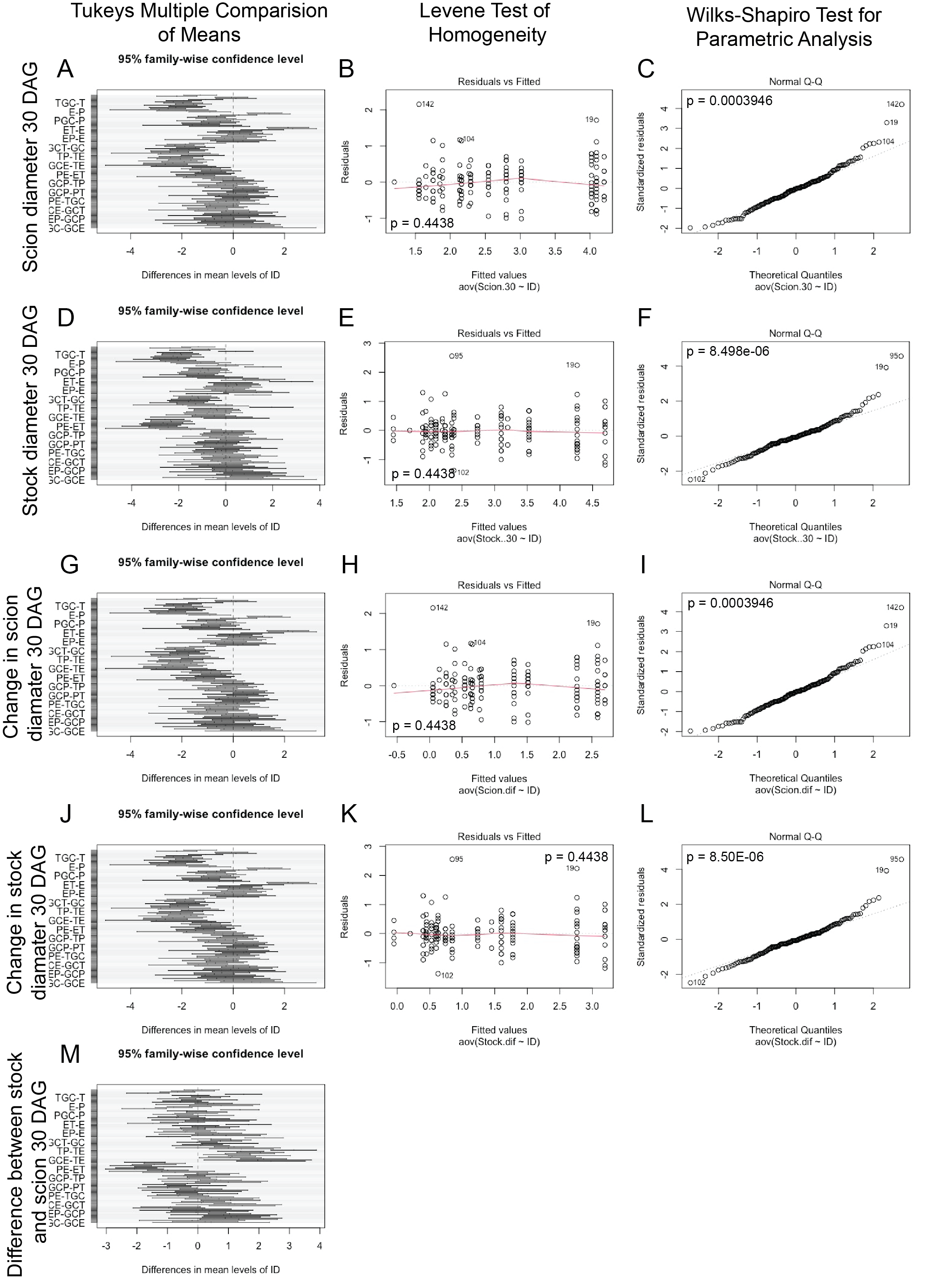

**Figure S4: Tukey's Multiple Comparison of Means, Levene's Test and Wilks-Shapiro Tests.** Tukey's Multiple Comparison of Means Test was used to perform pairwise comparisons (A, D, G, J, M). Levene’s Test of Homogeneity was used to determine if the data was homogenous (B, E, H, K). A p-value above 0.05 means the data is homogeneous. Wilks-Shapiro test was used to determine the parametric nature of the data (C, F, I, L). A p-value above 0.05 means the data is normal.
